## Appendix Table S5 for "Kinesin Autoinhibition Requires Elbow Phosphorylation"

**Appendix Table S5. *C. elegans* strains used in this study**

| **Strain name** | **Genotype** |
| --- | --- |
| *SYD0199* | *osm-3::gfp knock-in* |
| *PHX7125* | *syb7125[osm-3-pd::gfp knock-in]* |
| *PHX7130* | *syb7130[osm-3-pm::gfp knock-in]* |
| *GOU5378* | *syb7125[osm-3-pd::gfp knock-in]; syb7478[dyf-1::mScarlet knock-in]* |
| *GOU5379* | *syb7130[osm-3-pm::gfp knock-in]; syb7478[dyf-1::mScarlet knock-in]* |
| *GOU5380* | *cas11335[osm-3-pd + A489T::gfp]* |
| *GOU5381* | *cas11336[osm-3-pm + A489T::gfp]* |
| *GOU5382* | *osm-3(p802); casEx7031[*P*dyf-1::osm-3-pd + A489T::gfp; pRF4(+)]* |
| *GOU5383* | *osm-3(p802); casEx7032[*P*dyf-1::osm-3(T489E)::gfp; pRF4(+)]* |
| *GOU5384* | *osm-3(p802); casEx7033[*P*dyf-1:: osm-3(T489A)::gfp; pRF4(+)]* |
